# Population structure can reduce clonal interference when sexual reproduction and dispersal are synchronized

**DOI:** 10.1101/2023.07.10.548343

**Authors:** Qihan Liu, Daniel B. Weissman

## Abstract

In populations with limited recombination, clonal interference among beneficial mutations limits the maximum rate of adaptation. Spatial structure slows the spread of beneficial alleles; in purely asexual populations, this increases the amount of clonal interference. Beyond this extreme case, however, it is unclear how spatial structure and recombination interact to determine the amount of clonal interference. This interaction is particularly interesting because dispersal and recombination are often at least partially synchronized in natural populations, both at the individual and population levels, as when plants switch from vegetative growth to sexual reproduction or stress responses increase both motility and recombination in microbes. We simulate island models of populations evolving on a smooth fitness landscape and find that synchronized dispersal and sexual reproduction allow them to adapt faster than well-mixed populations of equal size. This is because the spatial structure preserves genetic diversity, while the synchronization increases the chance that recombination events occur between diverged individuals from different demes, i.e., the pairings where negative linkage disequilibrium can most effectively be reduced.

## INTRODUCTION

On smooth fitness landscapes, adaptation is driven by the fixation of beneficial mutations. When beneficial mutations are rare, they can fix independently from each other in sequential selective sweeps. But if the beneficial mutation supply is large, multiple beneficial mutations will be simultaneously polymorphic in the population, and may compete with each other for fixation. This “clonal interference” effectively puts an upper limit on the rate of adaptation, particularly when recombination is limited [Muller, 1932, Gerrish and Lenski, 1998, Desai and Fisher, 2007, Neher et al., 2010, Weissman and Barton, 2012]. Spatial structure increases the time it takes for a selective sweep [Slatkin, 1981, Whitlock, 2003] (in the absence of dominance [Sudbrack and Mullon, 2024]), and therefore increases the probability that multiple beneficial mutations will coexist and interfere. In asexual populations, strong spatial structure can drastically reduce the rate of adaptation [Martens and Hallatschek, 2011]. On rugged fitness landscapes, in contrast, populations must find the best combinations of mutations to adapt, and these combinations may not involve the mutations with the largest individual benefits. It has long been suggested that spatial structure may facilitate adaptation in this case [Wright, 1932, Kryazhimskiy et al., 2012, Nahum et al., 2015, Van Cleve and Weissman, 2015, Bitbol and Schwab, 2014, Covert III and Wilke, 2014, Cooper and Kerr, 2016, Crona, 2018], although it is controversial whether it actually does so in nature [Coyne et al., 1997].

To a certain extent, these two opposing effects of space on adaptation on smooth and rugged landscapes are caused by the same mechanism: spacial structure impedes competition and thus maintains genetic diversity. When an asexual population climbs up a smooth fitness peak, the rate of adaptation is always maximized by maximizing the reproduction of the fittest genotype currently present in the population; thus, spatial structure necessarily increases clonal interference and slows down adaptation [Martens and Hallatschek, 2011, Kryazhimskiy et al., 2012, Nahum et al., 2015]. However, the situation is less clear when recombination is introduced. At least in some situations, adaptation is primarily driven by recombination between genotypes that are not exceptionally fit. This can certainly be the case on rugged fitness landscapes [Wright, 1932, Cooper and Kerr, 2016, Crona, 2018], but it can also be true on smooth ones [Pearce and Fisher, 2019]. Thus while maximizing the reproduction of the fittest individuals maximizes the rate of adaptation in the current generation, it also reduces the genetic diversity that would fuel future adaptation [Robertson, 1970, Ueda et al., 2017]. This suggests that by slowing selection, spatial structure may actually be able to increase the rate of adaptation, even in smooth fitness landscapes.

Clonal interference is most likely in species in which individuals exchange genetic material only occasionally (as this limits recombination). This would include species that are only facultatively sexual or facultatively outcrossing, as well as asexual species such as bacteria in which recombination occurs via processes separate from reproduction. While the simplest model is that these occasions of genetic exchange are independent random occurrences, evidence suggests that they are often at least partially synchronized across individuals. Perhaps the most famous examples are of flowering synchrony and masting in plants [Kelly, 1994]. Many species of bamboo, for instance, alternate long periods of vegetative growth with bursts of synchronized sexual reproduction [Janzen, 1976]. Note that in plants, in addition to this synchrony of sexual reproduction at the population level, there is also a mechanistic synchronization at the individual level between sexual reproduction and dispersal.

While in some plants the bursts of dispersal and sexual reproduction occur in regular temporal cycles [Janzen, 1976], in other species, including many microbes, synchrony may be induced by the combination of fluctuating environmental conditions and the plasticity of dispersal and recombination in response to those conditions. In particular, both dispersal and recombination are often increased by stressful conditions (see examples reviewed in Gerber and Kokko [2018] and Rybnikov et al. [2023]). Mobile microbes can generally gain mobility from exposure to stressful environments such as starvation, temperature shift, and exposure to toxins [Son et al., 2015, Mitchell and Kogure, 2006], in extreme cases manifesting as collective swarming behavior [Belas et al., 1998, Daniels et al., 2004]. Stress has also been found to increase sexual reproduction in facultatively sexual microbes such as yeast [Freese et al., 1982, Bernstein and Johns, 1989, Davey, 1998, Neiman, 2011, Kassir et al., 1988, Mai and Breeden, 2000]. Considering recombination more generally, bacteria have also been found to undergo increased recombination as a general response to stress [Chung et al., 1989, Redfield, 1993, Lorenz and Wackernagel, 1994, Claverys et al., 2006].

The fact that dispersal and recombination are triggered by some similar environmental stimuli will tend to introduce some synchronization, both between the two processes (individual synchronization) and within processes across individuals (population synchronization, assuming that some environmental fluctuations affect many individuals in the population) [Iwasa and Satake, 2004, Gerber and Kokko, 2018]. It has been shown that population-level synchronization of facultative recombination can affect the rate of adaptation on both smooth [Weissman and Barton, 2012, Pearce and Fisher, 2019, Kokko, 2020, Berríos-Caro et al., 2021] and rugged landscapes [Weissman, 2014, Crona, 2018]. But these results are largely from well-mixed populations, and it is still unclear how synchronization of recombination interacts with population structure and the possible additional synchronization with dispersal. Intuitively, it seems plausible that population structure could allow diversity to build up in the population in different demes, which could then be brought together to produce new genotypes in a burst of synchronized dispersal and recombination, allowing the population to balance exploration and exploitation of the fitness landscape (Fig. 1).

**Figure 1.**
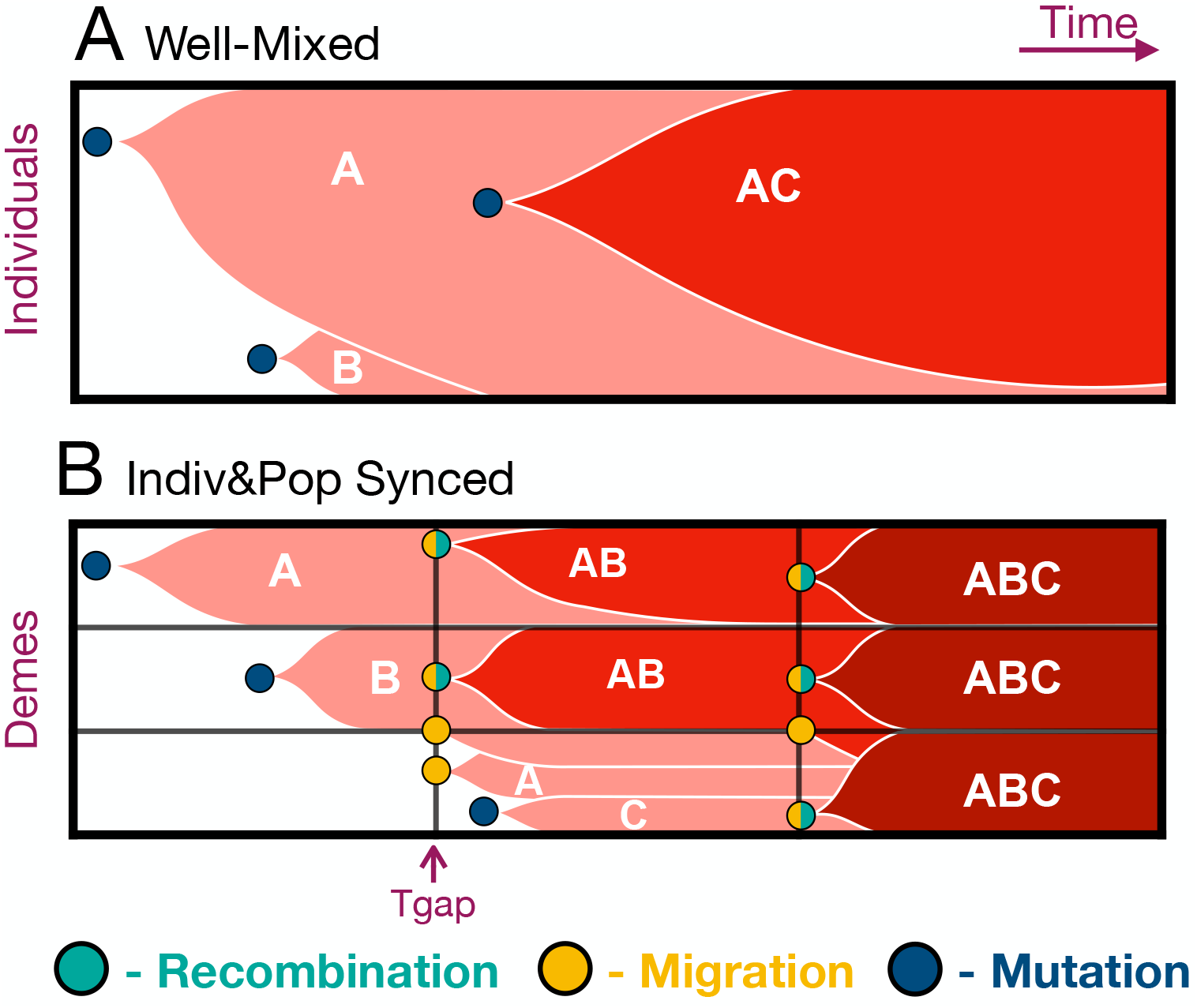
Schematic illustration of how population structure and synchronization can combine to accelerate adaptation. Darker red represents higher fitness, i.e., more mutations. **(A)** In a well-mixed population, mutations can rapidly spread to the whole population but drive all other diversity extinct in the process, with few opportunities for recombination between competing genotypes. **(B)** In a structured population, sweeps are slower but this preserves other beneficial mutations, which then can be brought together by synchronized dispersal and sexual reproduction, leading to faster long-term adaptation.

In this paper, we simulate evolution on smooth fitness landscapes and test the effects of population structure and fluctuating, possibly synchronized, recombination and dispersal on the rate of adaptation. We find that when the beneficial mutation supply is high, structured populations with individual- and/or population-synchronized recombination and dispersal can adapt faster than well-mixed ones. Population-level synchronization has the strongest effect, as it maximizes the probability that recombination events will occur between individuals with different beneficial mutations, the pairings that have the highest potential for generating extremely fit offspring. This process is most efficient when recombination happens simultaneously or shortly following dispersal.

## MODEL AND METHODS

Before we specify the details of the model, we will briefly describe the biological motivation for it which is drawn from yeast evolution experiments; see Fig. 2C of Kryazhimskiy et al. [2012] for a schematic of the kind of setup that we are modeling. Consider a population with cells grown in batch culture in many separate wells. Almost all division is asexual and takes place within the wells. One “generation” in the model corresponds to one transfer, i.e., the transition from one inoculum to the next inoculum. Each generation therefore consists of each individual producing a very large number of offspring which are then culled down by sampling to produce the next inoculum, very similar to the picture underlying the Wright-Fisher model. In rare special generations, dispersal and/or sexual reproduction take place just prior to sampling. For dispersal, some cells from all the wells are mixed together into a single migrant pool, and then redistributed as a portion of the inocula for all the wells in the sampling step. For sexual reproduction, a fraction of the cells are induced to mate and produce offspring. When both dispersal and sexual reproduction take place in the same generation, they are done simultaneously, i.e., sexual reproduction is induced while some of the cells are in the migrant pool, so that mating takes place within the migrant pool. Since we will be using a Wright-Fisher approach for computational efficiency, none of these within-generation processes will be simulated explicitly, but we hope that laying them out helps in understanding the effective model that we actually simulate.

**Figure 2.**
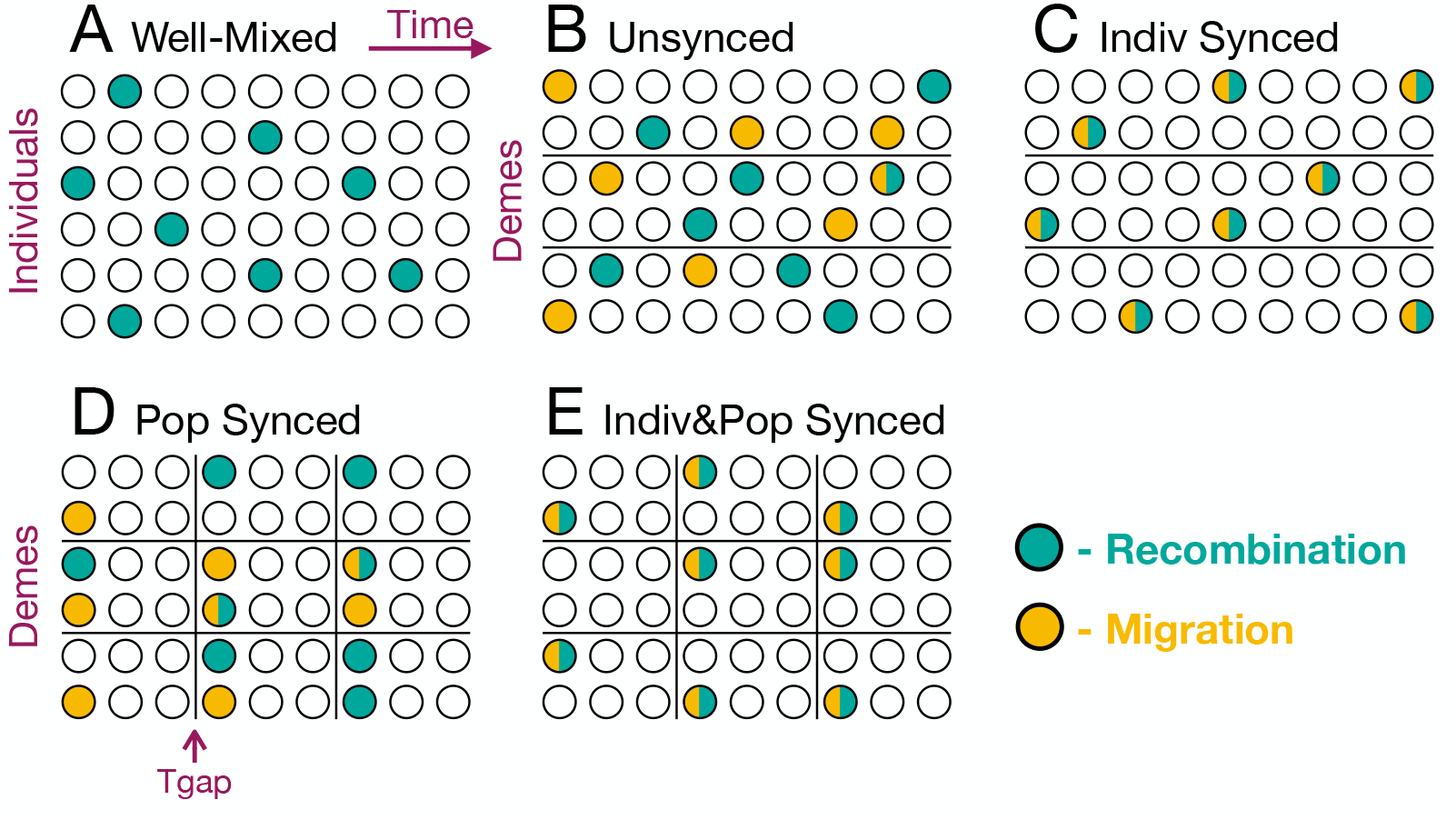
Illustrations of different possible forms of synchronization. **(A)** Well-mixed population with no synchronization of sexual reproduction. **(B)** Structured population (with deme boundaries shown as horizontal black lines) with no synchronization of dispersal or sexual reproduction. **(C)** Structured population with individual-level synchronization: different dispersal events are independent from each other, and similarly for instances of sexual reproduction, but dispersal and sexual reproduction are correlated with each other. **(D)** Structured population with population-level synchronization: dispersal and sexual reproduction are concentrated in generations that occur every *T*_gap_ generations (indicated by vertical black lines), but within one of these generations there is no association between the two processes. Population-level synchronization of sexual reproduction can also occur in well-mixed populations. **(E)** Structured population with both individual- and population-level synchronization: dispersal and sexual reproduction are correlated both between and within generations.

We consider a structured population distributed in *D* demes with *N* haploid individuals in each deme. We assume an island model, with all demes contributing equally to a single migrant pool and drawing from it at a time-dependent rate *m*(*t*) with average 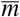. As a baseline, we also simulate a well-mixed population with only one deme with the same total population size, *D* × *N* individuals. Individuals are facultatively sexual, with outcrossing occurring at a time-dependent rate *f* (*t*) with average 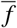. Each individual has *L* = 1000 unlinked loci. Beneficial mutations are uniformly distributed and occur at genomic rate *U*. We neglect back-mutations. Each mutation has the same log-fitness advantage *s* = 0.05 and all mutations combine multiplicatively, i.e., the fitness landscape is completely smooth. Because most reproduction is asexual, linkage disequilibrium can be large even though all loci are unlinked. We use this mating system and genetic map for simplicity, and do not believe that our results would be qualitatively different if we were to use a linear map or a model of gene conversion more similar to recombination in bacteria [Neher et al., 2010]. It remains to be seen what the quantitative effects of a more realistic mating system and map would be. Note that the number of simultaneously highly polymorphic loci is typically ≪ *L* in our simulations (see SI), suggesting that some quantitative results might persist even in models with loci arranged in a smaller number of chromosomes.

We allow for two possible ways that dispersal and recombination can be synchronized, as illustrated in Fig. 2. The first, individual-level synchronization, is a synchronization between the two processes and can be observed within a single generation. If the expected fractions of migrant and sexually produced offspring in that generation are *m*(*t*) and *f* (*t*) respectively, then there is individual-level synchronization if the expected fraction of offspring that are both migrants and sexually produced is *> m*(*t*) *f* (*t*). (See below for the precise details of how dispersal and reproduction are implemented in the simulations.) If, for example, *m*(*t*) *> f* (*t*), then in the most extreme case of individual-level synchronization, *all* sexually produced offspring would also be migrants. For simplicity, we focus on this extreme case in our simulations.

The other form of synchronization, population-level, can only be observed by comparing across multiple generations, and involves synchronizing dispersal, sexual reproduction, or both across individuals. It is automatically induced by the time dependence of *m*(*t*) and *f* (*t*). In generations when *m* is high, many individuals disperse, and in generations when it is low, few do; in generations when *f* is high, many individuals are the product of sexual reproduction, and in generations where it is low, few are. In populations with both forms of synchronization, both dispersal and sexual reproduction are concentrated in the same generations, and within those generations migrants are more likely than non-migrants to have been produced by sexual reproduction.

We consider a limiting form of population-level synchronization, in which dispersal and/or sexual reproduction occur only every *T*_gap_ generations, with *T*_gap_ = 1 corresponding to a steady rate, i.e., the absence of population synchronization. For *T*_gap_ *>* 1, then if the average probability of, e.g., dispersal is 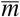, then there will be no dispersal for *T*_gap_ − 1 generations and then dispersal with probability 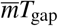 for a single generation. Larger values of *T*_gap_ therefore correspond to increased population-level synchronization. Note that since the probabilities of dispersal and sexual reproduction must both be ≤ 1 in each generation, 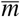 and 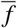 must both be less than 1*/T*_gap_. While this model is largely chosen for simplicity, it is also inspired by the design of microbial evolution experiments that test the effect of spatial structure [Kryazhimskiy et al., 2012, Nahum et al., 2015] or facultative outcrossing [McDonald et al., 2016] on the rate of adaptation. In these experiments, organisms have multiple generations of asexual reproduction within separate demes, interrupted by occasional generations of sexual reproduction and/or mixing among demes.

We use a form of Wright-Fisher reproduction, in which the entire population is replaced every generation. To produce an individual in deme *d* in generation *t* + 1, we first determine whether it is a resident (with probability 1 − *m*(*t*)) or a migrant (with probability *m*(*t*)). We then determine if it is the offspring of uniparental or biparental reproduction. In the absence of individual-level synchronization, these have probabilities 1 − *f* (*t*) and *f* (*t*), respectively. With individual-level synchronization, all resident individuals are produced uniparentally, while migrants have a probability *f* (*t*)*/m*(*t*) of being produced biparentally. (We always keep *f* (*t*) *< m*(*t*) in simulations with indivual-level synchronization.) Resident offspring draw their parent or parents from the individuals living in deme *d* at time *t* with probability proportional to their fitnesses. Migrant offspring draw their parents from a single pool comprising contributions from all demes. All demes contribute equally to the pool, but within each deme’s contribution, each potential parent’s contribution is weighted proportionally to its fitness. In other words, selection only acts within demes. (This is sometimes referred to as “soft” selection.) For migrants produced by biparental reproduction, the two parents’ demes are chosen independently, i.e., mating is assumed to take place within the pool. This setup is somewhat artificial for natural populations, but is perhaps closer to what might be implemented for experimental yeast populations [Kryazhimskiy et al., 2012], as described above. In the Results below, we show that a more standard model in which mating takes place after dispersal produces similar results.

The key outcome variable is the rate of adaptation *v*, defined as the rate of increase of mean log fitness. This is proportional to the probability of fixation of beneficial mutations, *P*_fix_: *v* = *NDU P*_fix_ ln(1 + *s*) ≈ *NDU P*_fix_*s*. In the absence of clonal interference, *P*_fix_ = 2*s* independent of the spatial structure [Maruyama, 1970], and the rate of adaptation is *v* = *v*_0_ ≈ 2*NDUs*^2^ [Weissman and Barton, 2012]. To understand the dynamics underlying the observed changes in *v*, we also track additional statistics of the populations. In particular, to determine whether adaptation is being driven by mutations in individuals already at the leading edge of the fitness distribution or recombinant offspring of parents in the bulk of the distribution leaping to the leading edge, we track the Hamming distance (the number of genetic differences) between the fittest individuals in the population in consecutive generations. See SI for additional statistics.

In our simulations, we keep *s* = 0.05 and *L* = 1000 for computational reasons— smaller values of *s* require longer run times, and larger values of *L* require more memory. We choose the other parameters to probe the region in which both selection and clonal interference are strong, 1*/N* ≪ *P*_fix_ ≪ 2*s*. To satisfy the first inequality, we primarily simulate large populations: all simulations use *N* = 10^6^ unless stated otherwise. The second inequality (strong clonal interference) requires that mutation be frequent and recombination rare. We determined that we could achieve this with computationally feasible total population sizes with mutation rate *U* = 5 × 10^−4^ per generation and average frequency of sexual reproduction 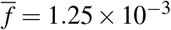 per generation. Unless otherwise stated, the number of demes in structured populations is *D* = 100, with average dispersal probability 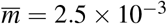, and the waiting time in population-synchronized simulations is *T*_gap_ = 100.

All plots show data averaged over 100 independent simulation runs, with error bars smaller than the size of the plot markers. In plots showing the average dynamics over the course of a cycle of *T*_gap_ generations, the first and last data points are the beginning of two consecutive bursty cycles, i.e., *T*_gap_ + 1 generations are shown.

## RESULTS

### Population structure can very slightly speed adaptation even without synchronization

In asexual populations, clonal interference is stronger in spatially structured populations than it is in well-mixed ones [Martens and Hallatschek, 2011, Kryazhimskiy et al., 2012]. Sexual reproduction reduces clonal interference in both spatially structured [Martens and Hallatschek, 2011] and well-mixed populations, but how does it change their relative rates of adaptation, i.e., the effect of spatial structure? To investigate this, we first simulate populations with facultative sex and varying degrees of spatial structure but no synchronization (schematic: Fig. 2B; results: Fig. 3 and Fig. S2B, orange lines).

**Figure 3.**
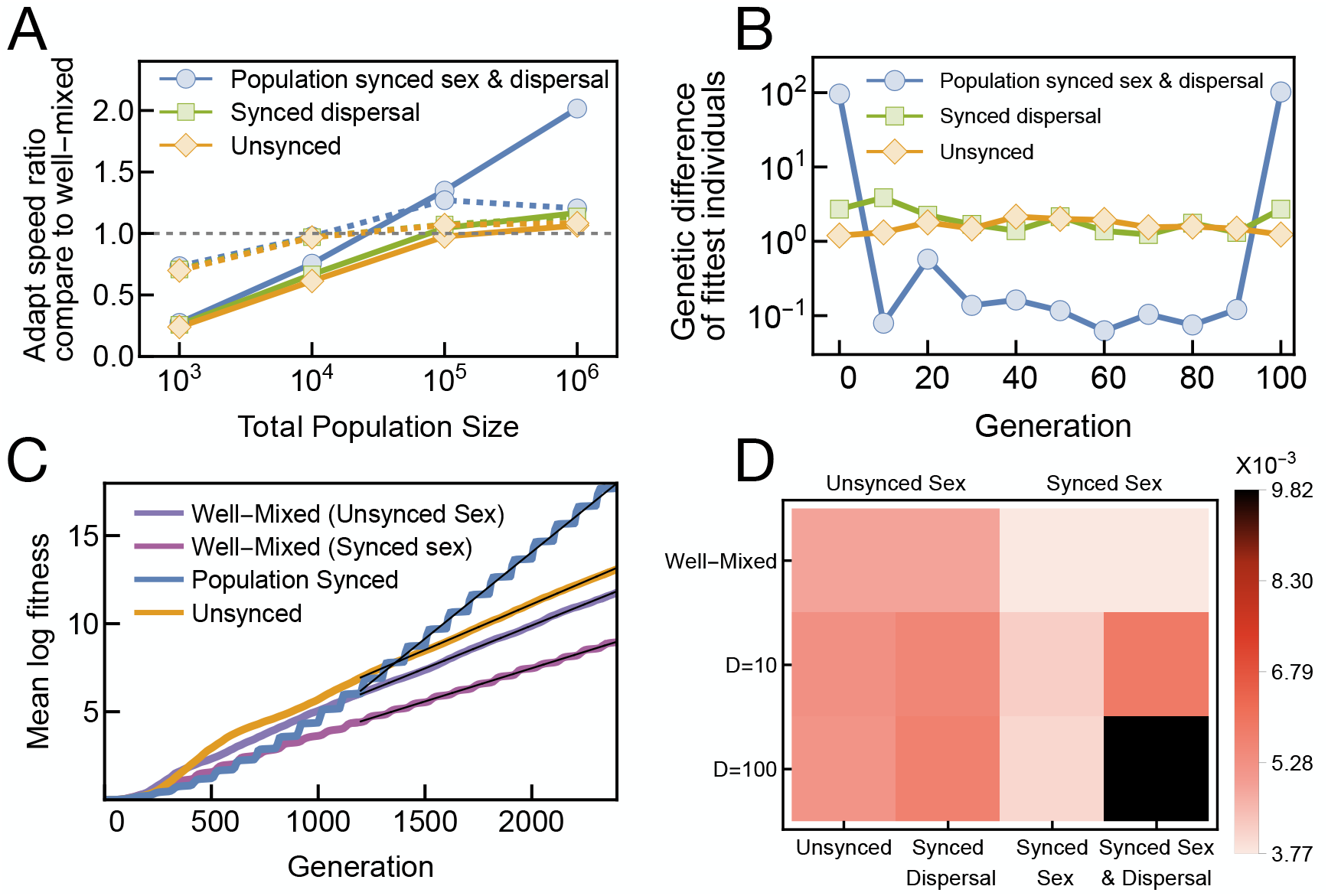
Population-level synchronization of dispersal and sexual reproduction accelerates adaptation by increasing the production of fit recombinants. **(A)** Ratios of adaptation speeds relative to a well-mixed population with unsynchronized sexual reproduction. Dashed lines show results from simulated populations comprising 10 demes, while solid lines show populations comprising 100 demes. Clonal interference is modest for the smallest populations of total size 10^3^ (adaptation is ∼ 3× slower than the maximum rate *v*_0_), increasing to very strong for the largest populations of total size 10^6^ (∼ 1000× slower than *v*_0_.) The strongly structured (100-deme) population with synchronized sexual reproduction and dispersal adapts roughly twice as fast as other populations at the largest simulated population size. Structured populations with no synchronization or synchronization of only dispersal have slight advantages over the well-mixed population (ratios 1.06, *p* = 0.035, and 1.17, *p <* 0.001, respectively, for populations of 100 demes). Details of the adaptation speeds are in Fig. S2. In all other simulation results, *D* = 100 and *N* = 10^6^ unless stated otherwise, since they produce the fastest adaptation. **(B)** Hamming distance between the fittest individuals in successive generations, shown over the course of the *T*_gap_ = 100 generations between rounds of synchronized dispersal/sexual reproduction. All curves are averages over 10 runs, each of which consists of 2400 generations (i.e., 24 *T*_gap_ periods). For populations without synchronization or synchronization of only dispersal, the small (≈ 1 mutation) distances indicate that adaptation is primarily driven by the accumulation of mutations on already-fit backgrounds. Populations with synchronized sexual reproduction and dispersal, on the other hand, leap ahead suddenly by ≈ 100 mutations due to recombination between divergent parents. **(C)** Mutation accumulation trajectories of different populations. The adaptation of the synchronized population is punctuated. Thin black lines indicate linear fits used for determining adaptation speeds. **(D)** Adaptation speeds (rates of increase of mean log fitness) of different populations. The full combination of strong structure and synchronized sexual reproduction and dispersal is needed to substantially accelerate adaptation. Individually, these factors only have very small effects or even slow down adaptation.

At total population sizes such that the mutation supply is limited but not tiny (*DNU* ≈ 1), clonal interference is just starting to affect well-mixed populations, and they still adapt at rates on the order of the maximum rate *v*_0_. (When the mutation supply is very low, *DNU* ≪ 1, lower than in our simulations, clonal interference is basically absent in well-mixed populations, and they adapt at close to the maximum rate even if they are asexual.) For our simulated parameters in Fig 3A, this corresponds to the smallest total population size, *DN* = 10^3^ = 0.5*/U*, for which well-mixed population adapts at ≈ 40% of the maximum rate of adaptation, *v*_0_ = 2.5 × 10^−3^ (Fig. S2). In this regime, increasing spatial structure only impedes adaptation (i.e., the left points in Fig 3A are all below one), because it slows down sweeps, increasing the probability that they overlap and interfere with each other. However, for large mutation supplies (*DNU* ≫ 1), well-mixed populations with limited recombination experience strong clonal interference. Surprisingly, in this regime, splitting the population into demes does not slow down adaptation and even very slightly accelerates it, with the population split into 100 demes evolving 6% faster than the well-mixed population (t-test, *p* = 0.035; see Fig. 3A, right side).

To understand this result, note that under strong clonal interference the speed of adaptation is only weakly (logarithmically) dependent on population size [Desai and Fisher, 2007, Neher et al., 2010, Weissman and Barton, 2012, Weissman and Hallatschek, 2014]. Thus, in a population large enough that each individual deme has a high mutation supply, each deme can adapt nearly as fast as a well-mixed population of the same size as the whole meta-population. So space only slightly slows down asexual evolution, and the increased genetic diversity between demes means that spatially structured populations can benefit more from recombination, pushing their overall rate of adaptation above well-mixed ones (Figs. 3A and S1).

### Population synchronization of sexual reproduction and dispersal can significantly accelerate adaptation

We first examine the effects of pure population synchronization, in which both sexual reproduction and dispersal are synchronized at the population level, with no additional individual-level synchronization (schematic: Fig. 2D). This represents a scenario in which, for example, an entire experimental population is forced to go through periodic rounds of dispersal and sexual reproduction [Kryazhimskiy et al., 2012, McDonald et al., 2016]. We see that when clonal interference is strong, this life history can approximately double the speed of adaptation relative to that of a well-mixed population with no synchronization (Fig. 3A). The increase in speed relative to a well-mixed population with the same synchronization of sexual reproduction is even larger (Fig. S2A). Thus, spatial structure reduces clonal interference in this instance. Spatial structure and synchronization act synergistically: in the absence of synchronization, spatial structure only accelerates adaptation by a tiny amount, while in absence of spatial structure and synchronized dispersal, synchronization of sexual reproduction actually slows down adaptation (Fig. 3D and Fig. S2).

The population with synchronized sexual reproduction and dispersal achieves its higher adaptation speed via bursts of adaptation every *T*_gap_ generations. These bursts move both the nose and the mean of the fitness distribution. But the nose (Fig. 3B) shifts in genotype space by far more than the distribution moves toward the optimal genotype (Fig. 3C). This indicates that the leading genotype is replaced by a distantly related recombinant, unlike in the other populations. The combination of structure and synchronization apparently allows it to both maintain and exploit a large reservoir of beneficial mutations beyond those that are found in the current best genotype.

### Sexual reproduction among migrants is crucial to the increase in adaptation speed

We next introduce individual-level synchronization. Such synchronization could arise as a response to individual-level environmental variation, as opposed to the global environmental fluctuations that would lead to population synchronization. We find that it can also accelerate adaptation, although not by as much as population synchronization (Fig. 4A, yellow squares vs dark blue diamonds). Populations with both forms of synchronization adapt fastest of all (Fig. 4A, light blue triangles), although the increase in speed over pure population-level synchronization is modest at the large population sizes where synchronization accelerates adaptation the most.

**Figure 4.**
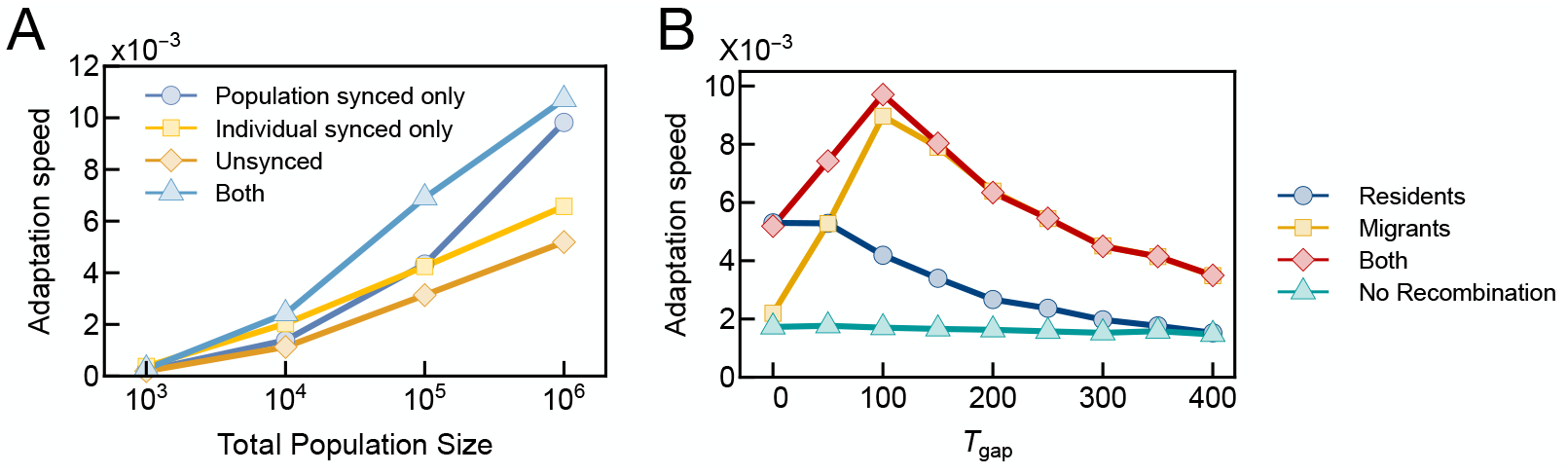
Sexual reproduction among migrants drives the increase in adaptation speed. **(A)** Adaptation speed of populations with differing synchrony between sexual reproduction and dispersal. Individual-level synchronization between sexual reproduction and dispersal accelerates adaptation, although not by as much as population-level synchronization. The two forms of synchronization act synergistically at moderate population sizes but have diminishing returns at larger population sizes. **(B)** Effect on adaptation speed of limiting sexual reproduction to migrant or resident individuals in populations with population-synchronized sexual reproduction and dispersal. For low values of the synchronization parameter *T*_gap_, almost all individuals are residents and sexual reproduction among migrants contributes negligibly to adaptation. At higher values of *T*_gap_ ≈ 100, even though migrants are still a minority, it is their offspring who are driving adaptation, with the majority residents making a negligible contribution. For even higher values *T*_gap_ *>* 200, migrants make up a majority of the population in the generations in which sexual reproduction takes place.

There are two possible explanations for why individual-level synchronization provides only a modest acceleration in large populations that already have population-level synchronization: either it is not important that sexual reproduction be happening specifically among migrants, or population-level synchronization already creates a large enough pool of sexually reproducing migrants that there are only limited benefits to increasing it further. To test these possibilities, we modify the simulations with only population-level synchronization so that either only migrants or only residents can reproduce sexually. We do not increase the rate of sexual reproduction among the group where it is allowed, so this lowers the overall rate of sexual reproduction—drastically so when sexual reproduction is limited to migrants, who are typically a minority of the population. For example, at the value *T*_gap_ = 100 where synchronization provides the greatest benefit, 25% of individuals are migrants in the high-dispersal generations. But we see that limiting sexual reproduction to these migrants hardly slows down adaptation, while limiting it to the 75% of individuals who are residents reduces the rate of adaptation by more than a factor of two (Fig. 4B). We therefore see that sexual reproduction among migrants is essential to the advantage of synchronization, suggesting that the limited benefits of individual-level synchronization are simply because population-level synchronization already creates a strong association between the two processes. It makes sense that sexual reproduction among migrants is the most important, as they are likely to differ by the most genetically and therefore have the most potential for producing extremely fit offspring.

### The synchronization *between* sexual reproduction and dispersal is crucial to the increase in adaption speed

Population-level synchronization both synchronizes dispersal with sexual reproduction and synchronizes among multiple dispersal events and among multiple instances of sexual reproduction. It seems intuitive that the former effect should be the most important for adaptation, since it focuses sexual reproduction on pairings with the greatest genetic diversity. But individual-level synchronization also produces this effect, and does not accelerate adaptation nearly as much (Fig. 4A). This raises the question of whether perhaps the synchronization between sexual reproduction and dispersal is actually unimportant, with the acceleration being driven by just the separate synchronizations among dispersal events and among instances of sexual reproduction. To test this, we introduce an offset between generations of increased dispersal and generations of increased sexual reproduction. Experimentally, this would correspond to having separate passages in which cells were mixed across wells and in which some cells were forced to undergo sexual reproduction. In natural populations, it would correspond to different pathways triggering dispersal and sexual reproduction in response to different environmental cues.

We find that the rate of adaptation is maximized when sexual reproduction and dispersal occur in the same generation (Fig. 5A), confirming the initial intuitive expectation. The rate of adaptation decreases as the number of generations elapsing after dispersal and before sexual reproduction increases, as the genetic diversity introduced into demes by dispersal is lost. Reassuringly, a delay of 10 generations, which is already on the order of the selection time scale 1*/s*, only introduces a 20% reduction in the adaptation speed. Notice that while for offset 0 we assume that sexual reproduction involving migrants is taking place within the migrant pool, for offset 10 sexual reproduction is taking place within demes. The similarity in outcomes shows that our results do not depend on the unusual setup of our model, in which there is a separate pool of migrants with sexual reproduction taking place only in that pool—the same effects occur in a model in which sexual reproduction occurs within demes after dispersal.

**Figure 5.**
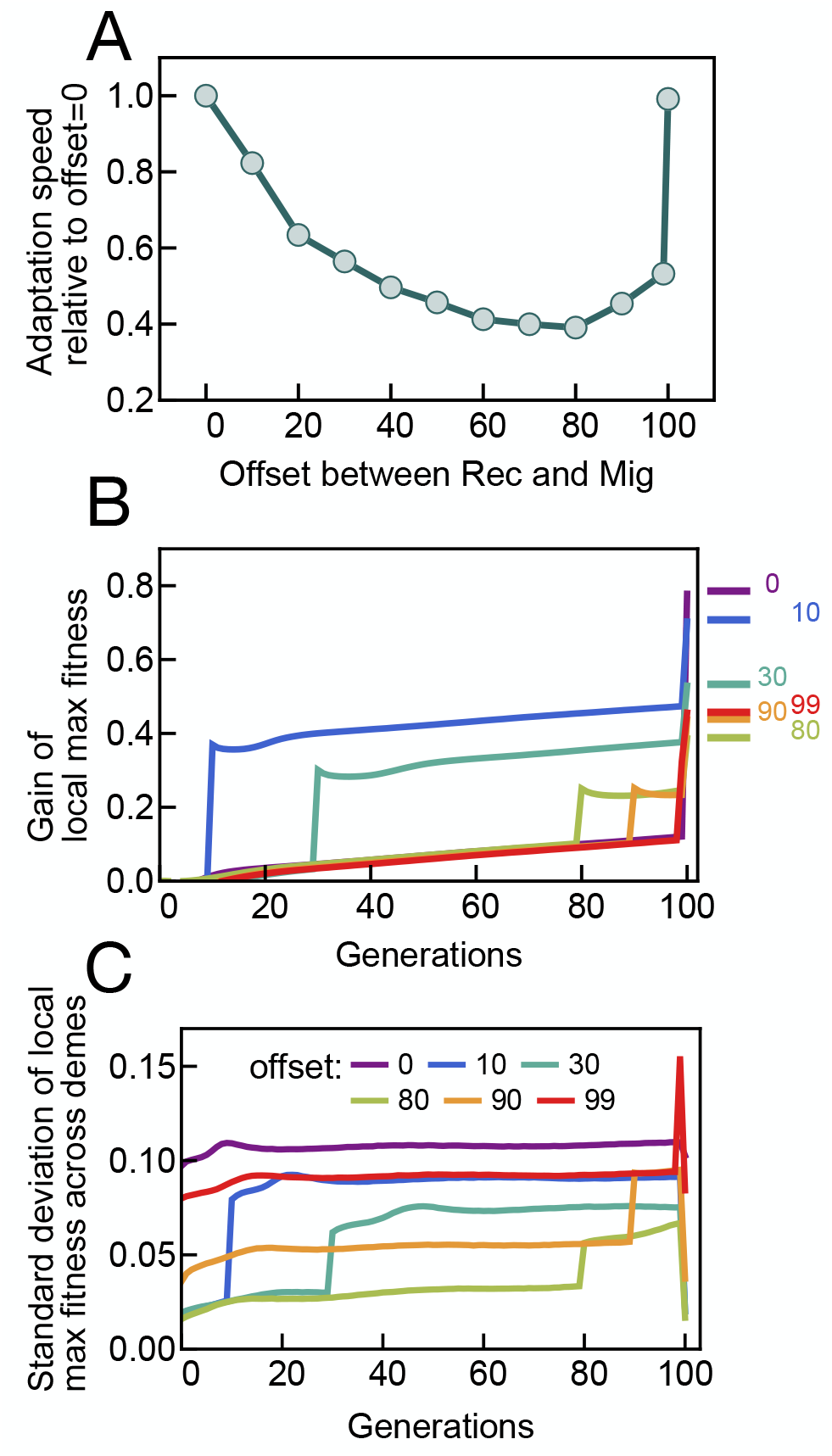
Simultaneous sexual reproduction and dispersal maximizes the rate of adaptation. **(A)** Adaptation speed as a function of the delay between high-dispersal generations and high-sexual reproduction generations. Dispersal occurs at times 0 and 100 on the horizontal axis, so an offset of 100 is identical to an offset of 0 and is only reproduced to highlight the contrast with an offset of 99. Increasing the delay before sexual reproduction lowers the rate of adaptation, as the increased within-deme genetic diversity introduced by dispersal is lost. Surprisingly, there is a slight uptick in the rate of adaptation for very long delays such that dispersal follows shortly *after* sexual reproduction. **(B)** Increase in the fitness of the fittest individual in a deme over the *T*_gap_ = 100 generations between dispersal events, for different values of the delay before sexual reproduction. This local maximum fitness jumps in the sexual reproduction generation, but by less as the delay increases and the genetic diversity introduced by dispersal decays. It appears to reach a steady state at ≈ 80 generations, partially explaining the uptick in panel A. **(C)** Standard deviation of local maximum fitness across demes shows the same qualitative pattern as overall adaptation speed (A), initially decreasing as the delay increases and then rebounding for very long delays. In all panels, points and curves are the averages over all the *T*_gap_ intervals in full simulation runs.

However, when we look at very long delays, such that dispersal quickly *follows* sexual reproduction, we find a new surprise: the rate of adaptation partially recovers (uptick at offsets of 90 and 99 in Fig. 5A). Examining the statistics of the simulations further reveals that the initial decrease in the rate of adaptation can be explained by a decrease in the fitness of the fittest recombinants formed (Fig. 5B). This effect saturates for long delays, likely because within-deme genetic diversity reaches a steady state maintained by mutation. At this point, it is better for adaptation to delay sexual reproduction more so that fit recombinants can quickly be redistributed across demes in the *next* dispersal event. This is because the demes vary in fitness (Fig. 5C), so dispersal will tend to move very fit recombinants into less-fit demes, where they will compete with each other less. For these very long offsets, the absolute adaptation speed (5.23 ± 0.11 × 10^−3^ for offset = 99) is similar to that for populations with synchronized dispersal but unsynchronized sexual reproduction (5.68 ± 0.15 × 10^−3^, Fig. S2C) or no synchronization (5.19 ± 0.12 × 10^−3^, Fig. S2B).

## DISCUSSION

In this paper, we demonstrate that population structure, by creating the possibility for synchronized dispersal and sexual reproduction, can increase the rate of adaptive evolution. To our knowledge, this is the first demonstration that a population structure that does not affect the fixation probability of isolated alleles can accelerate adaptation on smooth fitness landscapes without epistasis by reducing clonal interference among alleles. It does this both by alternately accumulating diversity in different demes and then bringing it together and forming fit recombinants, and by tending to concentrate recombination events in pairs of migrant individuals who are likely to be genetically distinct.

### Unexplored regions of parameter space and possible extensions

While the effects found here are substantial (e.g., a two-fold increase in adaptation speed relative to a well-mixed population), they are not enormous. This is in contrast to the situation on rugged fitness landscapes, where it has been suggested that structure and synchronized dispersal and recombination can have huge effects on the rate of adaptation [Crona, 2018]. One possibility is that we have not seen very large effects simply because we have not been able to simulate large enough populations. But our simulations do not suggest that much larger effects on the horizon, as the increase in the advantage of structure with population size appears to be modest, perhaps logarithmic (Fig. 3A, blue curve). A simple combinatorial argument also suggests that the gains should be limited: if, because of strong clonal interference, each of the *D* demes adapts nearly as fast as a well-mixed population of size *N* would during the *T*_gap_ generations between mixing, then the best-case scenario would be that each deme acquires unique mutations and recombination is frequent enough to break down linkage disequilibrium among all of them in the mixing generation. This would create a speed-up by a factor of nearly *D*. But the number of genotypic combinations that need to be explored by recombination during the mixing generation grows exponentially with *D*, so for any reasonable population size it seems more likely that the linkage disequilibrium created by the population structure will partially persist and the realized speed-up will be much less.

Besides considering larger values of *N* and *D*, one could also consider the effects of varying the strength of selection *s*. But we expect that most natural populations will at least approximately obey a form of the diffusion scaling, where the dynamics are only sensitive to “dimensionless” compound parameters like *DNs* or 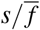. In other words, the dynamics should not be very sensitive to a rescaling of all parameters, and so one can choose to fix one of them (in our case, *s*) and then measure all other parameters relative to it. We have thus used a relatively “high” value of *s* = 0.05 for computational reasons, to maximize the amount of adaptation that we see per simulated generation, while still being low enough to avoid pathologies that can occur when *s* = 𝒪(1). Note that the compound parameter *DNs* is not particularly high, with a maximum value of 5 × 10^4^ in our simulations. The more commonly considered *DN*_e_*s*, where *DN*_e_ is the (total) “effective population size” based on neutral pairwise genetic diversity, is lower still. Since we do not simulate neutral variation we do not have a precise estimate of *DN*_e_, but it should be less than the value estimated from the selected variation, *DN*_e,sel_ ≡ *DNv/v*_0_ [Weissman and Barton, 2012], which is ∼ 10^3^ in our simulations. These populations would therefore not be considered particularly large if measured by the standard methods used for natural populations.

While we have observed that there is an intermediate value of *T*_gap_ that gives the strongest effects (Fig. 4B), we do not understand exactly what sets this value. Additional statistics discussed in the SI suggest that it is related to the time needed for selection to use up the burst in within-deme variation created by dispersal and recombination, but also to the time needed for new mutations to create inter-deme variation. Even if exploring a wider range of simulation parameter values would not greatly the maximum amount of acceleration, it could help clarify the scaling of the optimal *T*_gap_, which would help give more mechanistic insight into the observed effect.

Many of our modeling choices were made simply for convenience and simplicity, and alternate choices could be explored. As mentioned above, some forms of epistatic interactions are likely to have very large effects [Cooper and Kerr, 2016, Crona, 2018]. Extending the model to diploids, dominance might also be important. The selective coefficients of the mutations could be drawn from a distribution. As for recombination, as mentioned earlier, a more realistic model of either crossovers or homologous recombination could be used. Finally, there are wide range of possibilities for how to model synchronization. One obvious extension would be to include the evolution of synchronization within the model itself by adding modifier loci.

### Synchronization of dispersal and sexual reproduction in experiments and natural populations

Our model is meant to match the most straightforward way of implementing population structure and sexual reproduction in experimental yeast population grown in batch culture. Since dispersal is naturally done at transfers, it is automatically synchronized [Kryazhimskiy et al., 2012, Nahum et al., 2015]. Synchronized sexual reproduction (or more generally, recombination) is also natural when working with organisms such as yeast or many bacteria where it must be induced by environmental conditions [Chung et al., 1989, McDonald et al., 2016].

We believe that our model also captures important features found in natural populations. Facultative sexual reproduction is common in nature [Otto, 2009]. Sexual reproduction often carries costs, including mate-finding, performing sexual behavior, and producing males [Smith, 1978, Partridge and Hurst, 1998]. Facultative sexual reproduction provides most of benefits in terms of creating genetic diversity with less cost than obligate sexual reproduction [Kokko, 2020, Otto, 2009, Burke and Bonduriansky, 2017]. If sexual reproduction is rare, having it be synchronized helps reduce mate-finding costs [Kokko, 2020]. And dispersal is often linked to sexual reproduction (or outcrossing) in facultatively sexual populations, particularly in plants where there is often a mechanistic connection between them.

Facultative sexual reproduction is often a response to stressful conditions [Otto, 2009]. To the extent that these stressful conditions affect multiple individuals, this provides one mechanism for synchronization. As dispersal is also often a response to stress (e.g., [Martorell and Martínez-López, 2014, Gerber and Kokko, 2018, Kokko, 2020]), it will naturally also be synchronized with sexual reproduction. (Note that this condition dependence would also tend to create negative correlations between fitness and sexual reproduction and dispersal, but here we have assumed that this effect is small compared to the effect of non-genetic fluctuations [Weissman, 2014].) While environmental fluctuations are implicitly one possible source of synchronization in our model, the fitness landscape is static. Presumably the primary reason that sexual reproduction and dispersal can be triggered by stress is that it is a signal that the organism is poorly adapted to its present habitat, and that it should try to improve the match by producing offspring with different haplotypes or moving to a new environment. In other words, the response most likely evolved as a way to track the fluctuating components of the fitness landscape. Our work shows that a side effect can be more rapid adaptation to the fixed components of the landscape as well.

## ACKNOWLEDGMENTS

The authors thank Daniel Fisher for helpful discussions, and Julien Dutheil and two anonymous referees for their thorough and patient help in revising the work.

## CODE AVAILABILITY

All code for the simulations has been deposited on GitHub at https://github.com/weissmanlab/synchronized-sex-dispersal and is also available on Zenodo [Liu and Weissman, 2026].

## SUPPLEMENTARY INFORMATION

**Figure S1.**
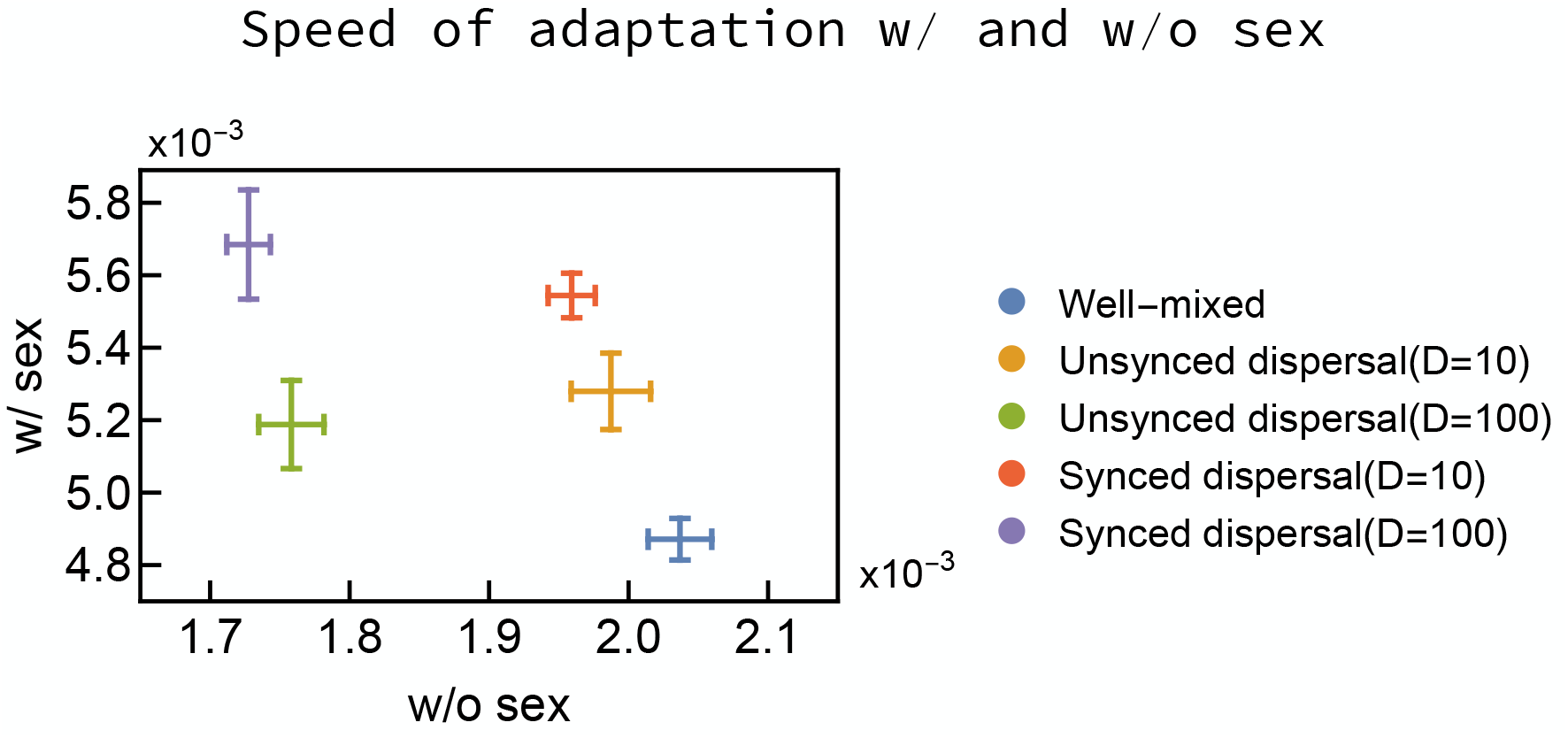
Structured populations benefit more from sexual reproduction than well-mixed populations. The vertical and horizontal axes are the adaptation speeds with and without sexual reproduction, respectively. Sexual reproduction is assumed to be unsynchronized. Without sexual reproduction, the well-mixed population adapts the fastest, while with sexual reproduction, it is the slowest.

**Figure S2.**
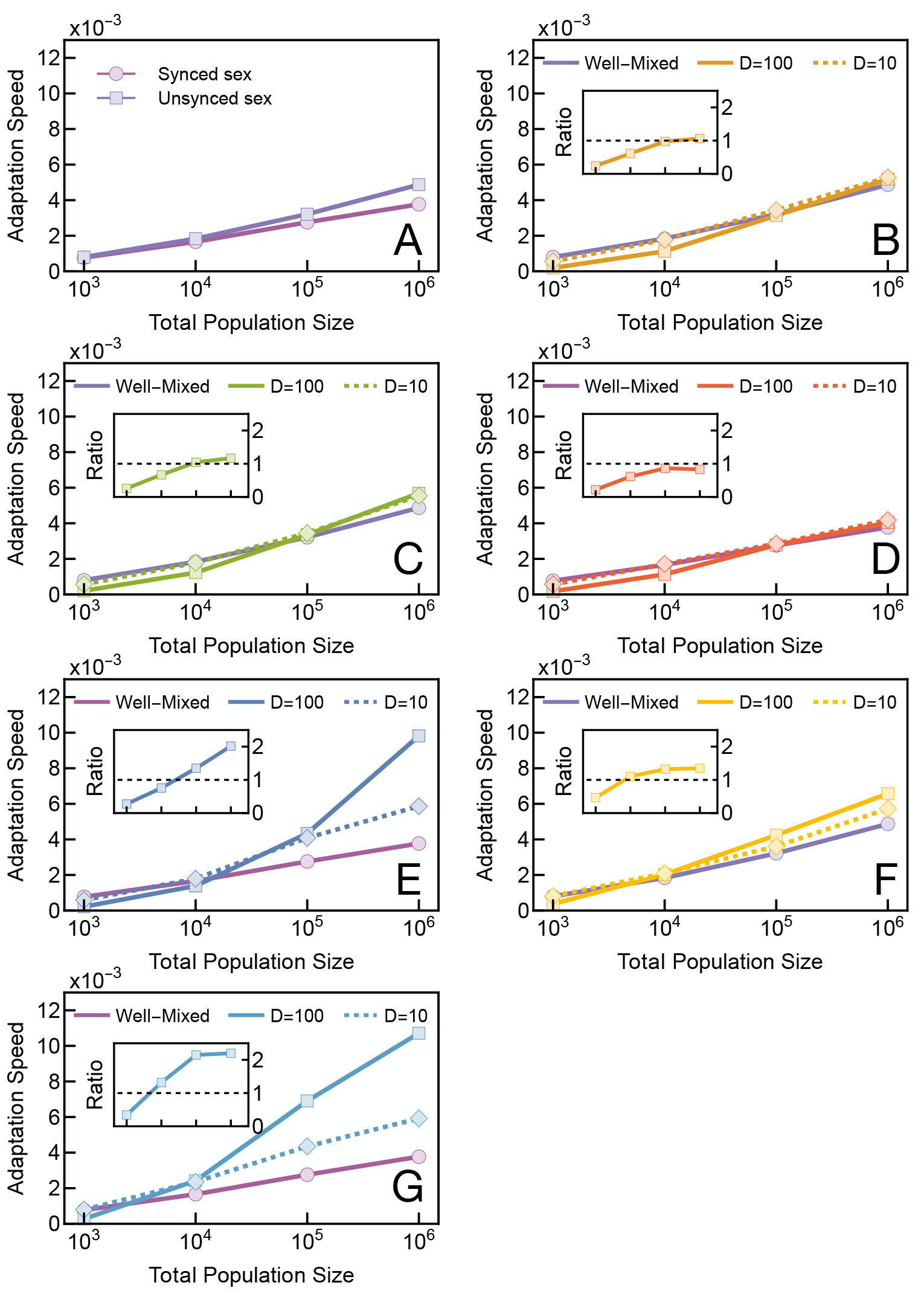
The adaptation speeds of different population structures. Insets show the ratio of the speed to that of a corresponding well-mixed population, which matches the focal population in whether its sexual reproduction is synchronized or unsynchronized. **A**. Well-mixed population. **B**. Unsynchronized structured population. All remaining panels show structured populations as well. **C**. Structured population with unsynchronized sexual reproduction and population-synchronized dispersal. **D**. Structured population with population-synchronized sexual reproduction and unsynchronized dispersal. **E**. Population synchronization for both sexual reproduction and dispersal with no additional individual synchronization. **F**. Individual synchronization with no population synchronization. **G**. Population and individual synchronization. **(B-G)**. The violet lines are the corresponding well-mixed population (with or without synchronization of sexual reproduction) from Fig. S2A. The solid lines are D=100, and the dashed lines are D=10.

### Genetic diversity and the nose of the fitness distribution

We track several additional population statistics in addition to those discussed in the main text. We measure total genetic diversity using the average pairwise genetic diversity 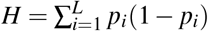, where *p*_*i*_ is the frequency of the mutant allele at locus *i*. If all *L* alleles were maximally polymorphic, *p*_*i*_ = 1*/*2, we would have *H* = *L/*4. 4*H* therefore serves as a measure of the effective number of highly polymorphic loci. Fig. S3 (panels A and C) shows that this is ∼ 10 for our largest well-mixed simulations, ∼ 30 when only dispersal is synchronized (∼ 20 in any given deme), and reaches a max of ∼ 200 (∼ 100 in any given deme) when dispersal and sexual reproduction are synchronized.

According to Fisher’s Fundamental Theorem [Fisher, 1930], the speed of adaptation is equal to the (genetic) variance in fitness. In our simulations, the standard deviation of log fitness is always ≤ 0.15 ≪ 1, which means that the variance in fitness is close to the variance in log fitness. In linkage equilibrium, this is simply *s*^2^*H*, proportional to the pairwise genetic diversity. The extent to which *v/s*^2^ lags behind the pairwise diversity *H* therefore provides a measure of total multilocus negative linkage disequilibrium. Comparing the results for *H* shown in Fig. S3 panels A and C to those for *v* in Fig. 3, we see that spatial structure increases genetic diversity by much more than it increases adaptation speed, i.e., it creates negative linkage disequilibrium. Strikingly, the genetic diversity in a single deme of the structured population can be higher than in the well-mixed population, even though its population size is 1% of the latter. Comparing Fig. S3 A and C to each other, we see that a similar pattern holds, with synchronization of sexual reproduction and dispersal increases global genetic diversity by a factor of ≈ 6 relative to only synchronized dispersal, more than the ≈ 2× increase in *v* (Fig. 3).

In addition to tracking the genetic differences between the fittest individuals from one generation to the next (Fig. 3B) over the cycle of *T*_gap_ generations, we also track their increase in fitness from the beginning of the cycle (Fig. S3, panels B and D). When only dispersal is synchronized (Fig. S3B), the fitness of the globally fittest individual increases steadily, while the average maximum fitness in each deme jumps up following the burst of dispersal in generation *T*_gap_, as individuals close to the global maximum fitness disperse into each deme. This then leads to a loss in global genetic diversity (dip in green curve in Fig. S3A) as demes converge on this type, before diversity is restored by new mutations. In contrast, when sexual reproduction is also synchronized (Fig. S3D), both the global and local maximum fitnesses jump up in generation *T*_gap_, as recombination between diverse individuals originating from differenting demes produces a burst of variation. This increase in global maximum fitness is ≈ 0.6, much less than the ≈ 100*s* ≈ 5 that would be the case if all the genetic differences (Fig. 3C) pointed in a consistent direction, suggesting the new fittest individual may be the offspring of parents that were both far from the previous fittest individual. Each deme will have its own unique fittest recombinants, so the demes do not converge genetically and global genetic diversity stays high (blue curve in Fig. S3C). Local genetic diversity decays in both sets of simulations.

**Figure S3.**
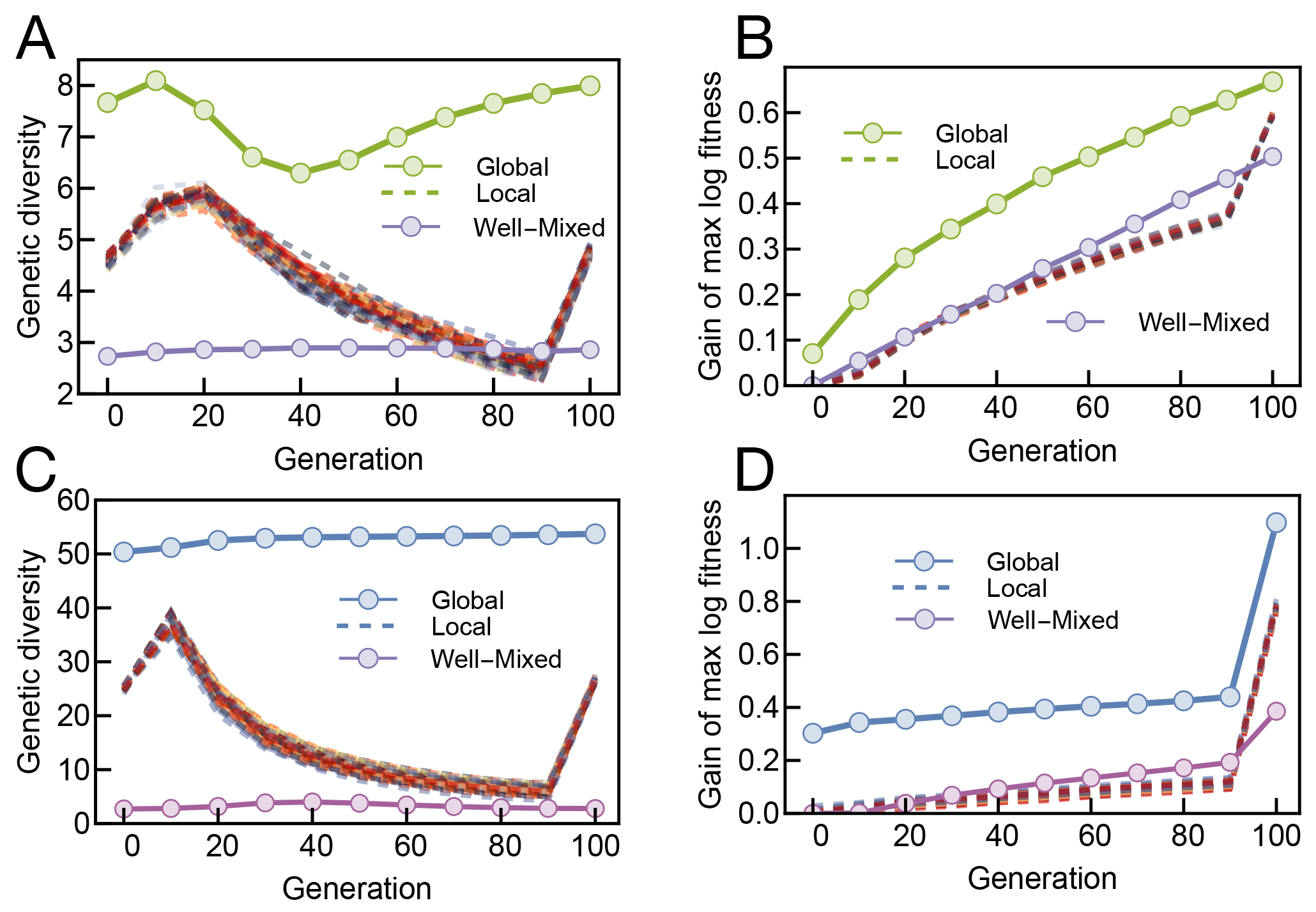
Synchronization of dispersal and sexual reproduction greatly increases genetic diversity and produces exceptionally fit recombinants. The panels contrast dynamics under population synchronization of only dispersal (A and B) and both sex and dispersal (C and D). Panels A and C show the pairwise genetic diversity *H* across the *T*_gap_ = 100 generations between events. Panels B and D show the average change in the log fitness of the fittest individual. The simulations and averaging procedure the same as in Fig. 3. All panels show statistics for the entire population, for each deme, and for a well-mixed population with the same synchronization or lack thereof of sexual reproduction.

